# Attenuation of HIV severity by slightly deleterious mutations can explain the long-term trajectory of virulence evolution

**DOI:** 10.1101/2025.05.12.653435

**Authors:** Harriet Longley, Christophe Fraser, Chris Wymant, Katrina Lythgoe

## Abstract

HIV-1 is a well-studied example of a pathogen that has evolved an intermediate level of virulence that maximises transmission. For a trait to evolve it must be heritable, and although viral load—a proxy for disease severity—has been shown to be a heritable trait, it is surprising that specific heritable viral factors remain mostly elusive. Rapid within-host evolution is also expected to diminish heritability. We hypothesised that rather than a small number of mutations of large effect determining viral load, the number of slightly deleterious mutations could be key. As a proof of principle, we explored how viral load is expected to evolve within and between hosts using a nested modelling approach that links within-host evolution with epidemiological outcomes. For mutations of sufficiently small effect, a mutation-selection balance is gradually reached during infection, resulting in slow changes in viral load despite rapid rates of genomic evolution. In simulated epidemics, we generated realistic population distributions of viral loads and estimates of heritability. The existence of many slightly deleterious mutations provides a mechanism that can help to explain why viral loads change slowly during infection, broad distributions of viral loads among individuals, and why searches for viral factors that determine viral load have had limited success.

## Introduction

HIV-1 has become a prominent example of a pathogen that has evolved towards intermediate virulence as a result of the trade-off between virulence—defined here as disease severity—and transmission (1). The trade-off hypothesis (2,3) is based on the idea that the duration of an infection and the virulence of the infecting pathogen are inversely correlated. If the relationship between virulence and transmission rate, saturates, the transmission fitness of the pathogen will be maximised at a finite level of virulence (4). If in addition virulence is heritable, meaning that differences in virulence between individuals is partly determined by genetic differences in the infecting pathogen, the pathogen is expected to evolve towards the level of virulence that maximises the number of onward transmissions.

Demonstrating the existence of evolutionary trade-offs of pathogens in real-world systems is notoriously difficult (4,5), and HIV-1 is one of the few systems where a transmission-virulence trade-off has been shown (1). During untreated chronic infection, HIV-1 viral loads typically remain at a relatively steady level, known as the set-point viral load (spVL). This is a striking observation given that spVLs are extremely heterogeneous when measured among individuals, varying by over four orders of magnitude (6). Because untreated individuals with higher viral loads tend to progress to AIDS and death sooner than those with lower viral loads (6,7), spVL is the most commonly used proxy for HIV-1 virulence. In addition, viral load has been shown to be correlated with infectiousness, with high viral load individuals much more likely to transmit the virus per contact (8,9). Fraser et al. (1) showed that the relationship between the expected number of onward transmissions and the duration of infection is saturating, and moreover that observed viral loads cluster around the values that are expected to maximise transmission, thus supporting the trade-off hypothesis. In a follow-up study, Blanquart et al. (10) found further support for the trade-off hypothesis from a large HIV-1 prospective cohort and argued that the attenuation of viral loads in the cohort over two decades was the result of between-host adaptation towards maximising transmission potential.

For a trait to evolve under natural selection it must be heritable. Although the study of heritability has its roots in animal breeding, the same concept can also be applied to pathogen virulence, i.e. for measuring the extent to which severity of infection is determined by the genotype of the infecting pathogen. Multiple studies have provided support for HIV-1 spVL being a heritable trait, with analyses proposing estimates of broad sense heritability ranging between 20-30% (11–17). However, the source of the heritability of HIV-1 spVL remains for the most part unexplained, suggesting that viral factors have small individual effect. Host-factors have also been attributed to variation in viral loads (18); however, a 2017 study found the fraction of variation explained by human genetic factors to be relatively low at 8.4% once viral genetic diversity is accounted for (19). Some human polymorphisms, such as the 32 base pair deletion in the CCR5 gene, have been found to significantly reduce HIV virulence (20,21). Virus-host interactions also likely affect virulence, for example the changes in immune escape pressure following transmission to a new host environment (22).

Another difficulty in understanding the heritability of HIV-1 spVL is rapid within-host evolution and often long delays between infection and onward transmission. HIV-1 has a high mutation rate per replication which, coupled with a short viral generation time, generates significant genetic diversity within individuals and the potential for rapid adaptive evolution (23). This process is likely to reduce heritability as the viral genotype an individual is infected with is expected to differ from the genotype that they go on to transmit (17,24). For example, we may expect viral variants with high replicative capacity to emerge and quickly sweep through the virus population, and if replicative capacity affects disease factors such as virulence, then the heritability of these factors will be reduced.

Viral replicative capacity has been shown to be associated with viral load (25–27). Even fairly small differences in the fitness of virus variants are expected to result in rapid within-host evolution, notwithstanding complications of whether the virus is pre-adapted to a recipient’s HLA allele type (28,29). Hence if there were a strong link between replicative capacity and viral load, we would expect viral loads to increase greatly during infection, low estimates of heritability, and the evolution of high virulence even at a cost to overall transmission (“short-sighted evolution”) (22,30). Yet viral loads only change modestly during chronic infection (31,32), heritability of viral load is remarkably high, and ever-increasing viral loads has not been a feature of the HIV-1 epidemic. Possible mechanisms for how virulence has evolved under the constraints of both within and between host selection include a complex fitness landscape, and the preferential transmission of ancestral strains (17). Underpinning any mechanism are the elusive heritable viral factors.

We propose that rather than being controlled by few high-impact mutations (as usually targeted by whole-genome association studies), viral load is largely determined by the number of slightly deleterious mutations a genome has. In such a scenario a mutation-selection balance, in which mutations continually arise through mutation and are slowly lost through purifying selection, is gradually approached during the course of infection. Because this process is inherently slow, it can reconcile seemingly incompatible aspects of the within- and between-host processes, namely the stability of viral load during chronic infection, the high heritability of viral load, and the selection of intermediate viral loads at the population level. Moreover, it may explain why HIV-1 virulence factors have been so hard to identify. As a proof of concept, we considered the impact of many slightly deleterious mutations on the within and between-host evolutionary dynamics of the virus within treatment-naïve individuals using a nested modelling approach.

## Methods

We modelled the evolving population of viral genotypes during infection using a quasispecies framework, with mutations occurring at a frequency of *μ* = 3 × 10^−5^ per site per generation (33) at *m* segregating sites. We assume each segregating site in the viral genome has two potential alleles, wild-type or slightly deleterious, with ‘virus types’ defined by the number of deleterious mutations in the viral genome, regardless of the position of those mutations, and hence viruses with different genotypes can have the same virus type. Individuals are assumed to be initially infected by a single virus type, which defines the infection type: an individual who was infected with virus type *j* will be of infection type *j.* After infection, the dual forces of a high mutation rate and weak selection result in the emergence of a diverse within-host viral population and the maintenance of deleterious mutations, with the population reaching mutation- selection balance. At time *t* during a type *j* infection, a virus type *i* has a frequency 𝑥_𝑖𝑗_(𝑡) that is determined by the quasispecies equation (see next sub-section). We assumed that the viral load of an infection at time *t* is determined by the number of deleterious mutations in the within-host viral population at time *t*. For the epidemiological (between-host) modelling, we use a nested modelling framework from Lythgoe *et al*. (30) in which the within-host dynamics are ‘nested’ within a between- host epidemiological model, which in turn is based upon the theory of multi-type epidemic models (34), and the course of an individual’s infection is entirely determined by the within-host model.

### Within-host dynamics

We assumed the viral genome has a maximum of *m* segregating sites, and a virus type is defined by the number of deleterious mutations, *j,* regardless of the position of the mutation. The generation time of an infected cell was assumed to be one day, and each newly infected cell can acquire or lose at most one mutation relative to the infecting virion. The *m x m* reproduction matrix 𝑄 = 𝑞_𝑖𝑗_ describes the probability that a progeny of virus type *j* is of virus type *i*, where 𝑞_𝑖𝑗_ is only non-zero when *i* and *j* differ by 0 or 1. We considered the competing virus types within a host as a quasispe (1) for which the dynamics are governed by the quasispecies equation (1), and the relative fitness of virus type *j* is a multiplicative function of the number of deleterious mutations: 𝐴_𝑗_ = (1 − 𝑠)^𝑗^ where *s* is the selection coefficient and 𝐴_0_ = 1. Here, viral fitness is synonymous with the replicative capacity of the virus type. The change over time in the relative frequencies of the virus types can be described by the quasispecies equation [38]:

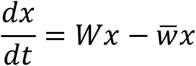

where 𝑊 = 𝑤_𝑖𝑗_ = 𝑞_𝑖𝑗_𝐴_𝑗_ and the term 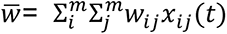 bounds the sum of the frequencies to one. The solution at equilibrium, 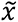, can be found analytically as the dominant eigenvector of *W* (35). The pre-equilibrium analytical solution of the system was solved numerically with the *deSolve* package in R v. 4.0.2.

The higher the fixed number of segregating sites in the model, the smaller the relative fitness cost of one additional mutation. We explored a range of fitness costs, from 10^−2^ per mutation when 10 sites are segregating to 5 × 10^−5^ per mutation when 250 sites are segregating, spanning strong selection to close to neutral effects approaching the mutation rate (36). We capped the maximum number of segregating sites, *m*, at 250 because the matrices needed to track all possible combinations of viral variants become too large to handle - they grow exponentially with each additional site we add to the model - however the range of values we considered effectively capture the key dynamics of the model.

### Viral load

There are three distinct stages of an HIV infection. The first and third stages of infection, termed acute and late infection respectively, are characterised by high viral loads. The second stage is chronic infection, when viral loads are relatively stable yet vary significantly between individuals. We assumed that during chronic infection the viral load at a given time is determined by the current composition of the viral population. As a consequence, the viral load can change as the viral population evolves: the fewer the number of deleterious mutations the higher the viral load. At any moment in time the viral population is likely to be comprised of multiple types, so we first define the contribution of each viral type, 𝑉_𝑖_, to the viral load during chronic infection:

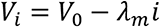

where 𝑉_0_ = 7 log_10_(viral copies per ml), *I* is the number of mutations, and 𝜆_𝑚_ is the reduction in viral load due to an additional mutation, which is in turn determined by the maximum number of mutations, *m*. The maximum possible viral load, 7 log_10_(viral copies per ml), was chosen to match the highest viral loads typically observed (17). The range was further verified in our dataset of spVLs.

When we assumed a large number of mutations, we assumed that the fitness cost of a single mutation is small, and therefore the reduction in viral load given one additional mutation is also small. Conversely, if we have few mutations of large effect, we assumed viral load will reduce significantly when a deleterious mutation appears. The value of 𝜆_𝑚_ is chosen such that the difference in the viral load between the fittest and weakest virus is 5 log_10_(viral copies per ml) to correspond with a realistic range of viral loads, and so 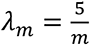. In determining viral load in this way, there is a linear relationship between the viral load - log_10_(viral copies per ml) - of a variant and the relative fitness of the variant. We then determined the realised viral load to be the mean of these contributions, so that viral load of a type *j* infection at time *t* into infection is given by:

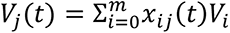

### Duration of chronic infection

We used the previously parameterised decreasing Hill function for the duration of chronic infection as a function of spVL (1). To account for the change in viral load over time in the calculation of the duration of infection, we took a weighted average of the viral load over the maximum possible duration of chronic infection, determined to be 20.4 years by the Hill function. The weight of 𝑉_𝑗_(𝑡) in the calculation of 𝑉̅_𝑗_ is inversely proportion to *t,* such that the viral load close to the start of chronic infection contributes more to the calculation than the viral load several years into infection. The weighted average viral load is then used as an input to the decreasing hill function to provide a chronic infection duration, termed 𝑇_𝑗_. To attribute a spVL, 𝑉_𝑗_, to an infection we take the arithmetic mean viral load over 𝑇_𝑗_.

### Infectivity profile

In order to determine how evolution proceeds at the between-host level, we need to know not only the frequency of different virus types during infection, but also their probability of transmission. We assumed a single virus type is transmitted during a transmission event, and the hazard for transmitting virus type *i* in a type *j* infection at time *t* is:

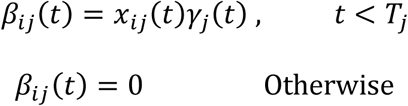

Where 𝛾_𝑗_(𝑡) is the infectiousness of a virus type *j* infection at time t and 𝑇_𝑗_ is the duration of a virus type *j* infection. For all infections we assume the acute phase has a duration of 0.25 years and an infectiousness of 2.76 onward infections/year, and the late phase has a duration of 0.75 years and an infectiousness of 0.76 onward infections/year (1). During chronic infection, the infectiousness 𝛾_𝑗_ (𝑡) is described by an increasing Hill function of viral load, *V(t),* at time *t* with parameter values based upon the optimal values detailed in (1).

**Table 1.** Variable and parameters descriptions for the within-host and between host models.

| Parameters | Definition | Value |
| --- | --- | --- |
| $\mu$ | Mutation rate | $3 \times 10^{-5}$ (33) |
| $m$ | The number of segregating sites, and therefore the maximum number of deleterious mutations. | 10, 50, 100, 150, 200, 250 |
| $s_m$ | The relative fitness cost of an additional mutation ranging from $s_{10} = 10^{-2}$ to $s_{250} = 5 \times 10^{-5}$ to capture a realistic distribution of selection coefficients. The values between $m=10$ and $m=250$ were determined by fitting a straight line between $s_{10}$ and $s_{250}$ on a log-linear scale to scale | $s_{10} = 10^{-2}$<br>$s_{50} = 10^{-2.4}$<br>$s_{100} = 10^{-2.9}$ |
| | selection strength with the number of segregating sites. | $s_{150} = 10^{-3.4}$<br>$s_{200} = 10^{-3.8}$<br>$s_{250} = 10^{-4.3}$ |
| $A_j$ | Replication rate of a type $j$ virus variant. | $A_j = (1 - s)^j$ |
| $\lambda_m$ | The cost of an additional mutation on the associated viral load. It is given by $\lambda_m = \frac{5}{m}$. The value is determined such that the minimum viral load is $2 \log_{10}$(viral copies per ml). | $\lambda_{10} = 0.5$<br>$\lambda_{50} = 0.1$<br>$\lambda_{100} = 0.05$<br>$\lambda_{150} = 0.033$<br>$\lambda_{200} = 0.025$<br>$\lambda_{250} = 0.02$ |
| $T_j$ | The duration of a type-$j$ infection. | Variable |
| $\gamma_j(t)$ | Infectivity of a type-$j$ infection at time $t$ into infection. | Variable and a function of $V_j(t)$. Parameterised in [1]. |
| $V_i^s$ | The viral load associated with a type-$i$ virus for a given fitness cost. | $7 \log_{10}(\text{copies per ml}) - \lambda_s i$ |
| $B$ | Rate at which individuals enter the susceptible population. | 200 per year (30) |
| $D$ | Natural mortality rate. | 0.02 per year (30) |
| $h$ | Host type. | 1-50 |
| $e$ | Maximum absolute host effect on viral load on $\log_{10}$ scale | 0.1, 0.25, 0.5, 0.75, 1 |
| $E_h$ | Additive host effect size on $\log_{10}$ viral load. | Discretely uniformly distributed over the range $[-e, e]$ in the host population. |
| Variables |  |  |
| $x_{ij}(t)$ | The frequency of a type-$i$ virus at time $t$ of a type-$j$ infection. | |
| $V_j(t)$ | Viral load at time $t$ of a type-$j$ infection | |
| $\beta_{ij}(t)$ | The infectivity profile i.e. the rate at which a type-$i$ virus is transmitted at time $t$ of a type-$j$ infection. | |
| $K$ | The next generation matrix. | |
| $I_i(t), I^*$ | The number of infected individuals infected with a type-$i$ virus at time $t$ into the epidemic ($I_i(t)$) and at the endemic equilibrium ($I^*$). | Initially 1 |
| $S(t), S^*$ | The number of susceptible individuals at time $t$ into the epidemic ($S(t)$) and at the endemic equilibrium ($S^*$). | Initially 10,000 |
| $N(t), N^*$ | The number of humans hosts at time $t$ into the epidemic ($N(t)$,) and at the endemic equilibrium ($N^*$). | Initially 10,000 |
| $h^2$ | Broad-sense heritability estimate. | |

### Between-host model

We used an SI model with demography to model between-host transmission, where we assumed a natural death rate *D* and that individuals enter the susceptible pool of individuals at a rate of *B*. The force of infection of virus type *i* at time 𝜏 of an infection founded by virus type *j* is defined as 𝛽_𝑖𝑗_ (𝜏)𝑒^−𝐷𝜏^when 𝜏 ≤ 𝑇_𝑗_ and 0 otherwise. The between-host dynamics are modelled by the renewal equation, as described in Lythgoe *et al.* (30). Specifically, incidence is calculated by the renewal equation which follows the logic that incidence at time *t* is the integral of the past incidences weighted by how much the individuals previously infected would still be transmitting. The between-host dynamics at time *t* since the beginning of the epidemic are therefore described as follows:

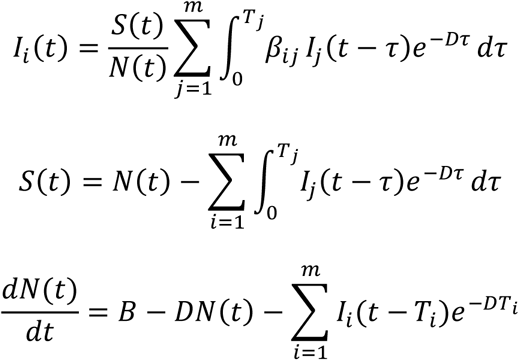

Where *N* is the total population size, *S* is the number of susceptible individuals and 𝑰_𝒊_ is the incidence of infections founded by a type *i* virus. All individuals have the same natural death hazard, *D*, and infected individuals also have an infinite death hazard at the moment their infection ends, which is pre-determined by their infection type.

We began the epidemic with an incidence of 1. The solutions were determined numerically using the basic forward Euler method. We ran 3 simulations of the between-host dynamics, where the initial circulating virus type and therefore average viral load differed for each simulation. Epidemiological theory shows that the next generation matrix *K*, with *ij*th element 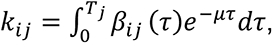, has a unique and real dominant eigenvalue that gives the value of the basic reproduction number *R_0_*, and the associated eigenvector describes the population structure of the virus types at equilibrium (30), which can be normalised to give the proportions of each virus type at their equilibrium state, 𝐼^∗^. The transmission potential of an infection by a type *j* virus is defined as the number of expected onward transmissions during infection and is determined from the next generation matrix as 𝑝 = Σ_𝑖_𝑘_𝑖𝑗_, where the term 𝑘_𝑖𝑗_is the expected number of transmissions of a type *i* virus during an infection of a type *j* virus.

### Host heterogeneity

As well as considering a population of genetically identical host individuals, we also examined the effect of host heterogeneity. The host effect is quantified by an additive effect to the viral load, such that for a type *j* infection in a host type *h*, the viral load at time *t* is given by 𝑉_j_^ℎ^(𝑡) = 𝑉_𝑗_(𝑡) + 𝐸_ℎ_. The parameter *E* is discretely uniformly distributed, 𝐸∼𝒰(−𝑒, 𝑒) and there are 50 possible host types. As a result, the probability of transmitting virus type *j* at time *t* differs by host type, however the within-host dynamics are identical for all infections. We considered a range of values of *e* from 0.1 to 1 in order to quantify the impact of host specific viral load effects on model outcomes. We present the analytical equilibrium solution for populations with host- heterogeneity as it was not feasible to derive pre-equilibrium solutions numerically due to computational constraints.

### Heritability

To estimate broad-sense heritability, ℎ^2^ (heritability hereafter for brevity), we use the classic parent-offspring regression method (37). To apply the method to a simulated infection population, we generated 1000 samples of 500 transmission pairs, where the source infection type was sampled based upon the population distribution of infection types at the endemic steady state. The probability of a recipient having an infection of a type *i* virus from a source infected with type *j* virus was described by the vector of transmission potentials, 𝑘_𝑖𝑗_, normalised to sum to 1. For each set of transmission pairs, an estimate of ℎ^2^ was given by the regression coefficient of a simple linear regression with recipient spVL as the outcome. The overall estimate of heritability is calculated by the average ℎ^2^ across the sets of transmission pairs. For a non-homogeneous host population, the host type of the source and recipient was also sampled from a uniform distribution of host types.

## Results

To summarise our approach, I modelled the within-host dynamics with a quasispecies model that describes the changing frequencies of competing virus types within a host virus population, where each virus type is defined by its number of deleterious mutations. The relative fitness of a virus type is a function of the number of deleterious mutations, and the conflicting forces of a high mutation rate and purifying selection results in a mutation-selection balance of deleterious mutations if the fitness cost is sufficiently low. The viral load at time *t* is determined by the average fitness of the virus population at time *t,* where high fitness induces a high viral load. The within-host model is nested in a between-host model that can describe the population-level distributions of viral loads and the amount of heritability for different parameter choices.

### Many mutations with low fitness cost results in intermediate virulence evolving within individuals

We determined the within-host equilibrium solution for a range of values for the number of segregating sites, *m*, and calculated the corresponding viral load (Figure 1). We find that for a relatively low number of sites (fewer than approximately 100), selection of the fittest virus types dominates, and viral load is at the maximum possible value. As the number of sites increases, the selection cost decreases, and we approach a mutation-selection balance. Consequently, the number of deleterious mutations in the population is maintained and the viral load is reduced. Eventually, the cost of an additional mutation passes a threshold where the relative fitness cost is effectively neutral and the equilibrium viral load approaches the midpoint of the viral load range of 4.5 log_10_(viral copies per ml), and the genetic variation is maintained through mutation.

**Figure 1:**
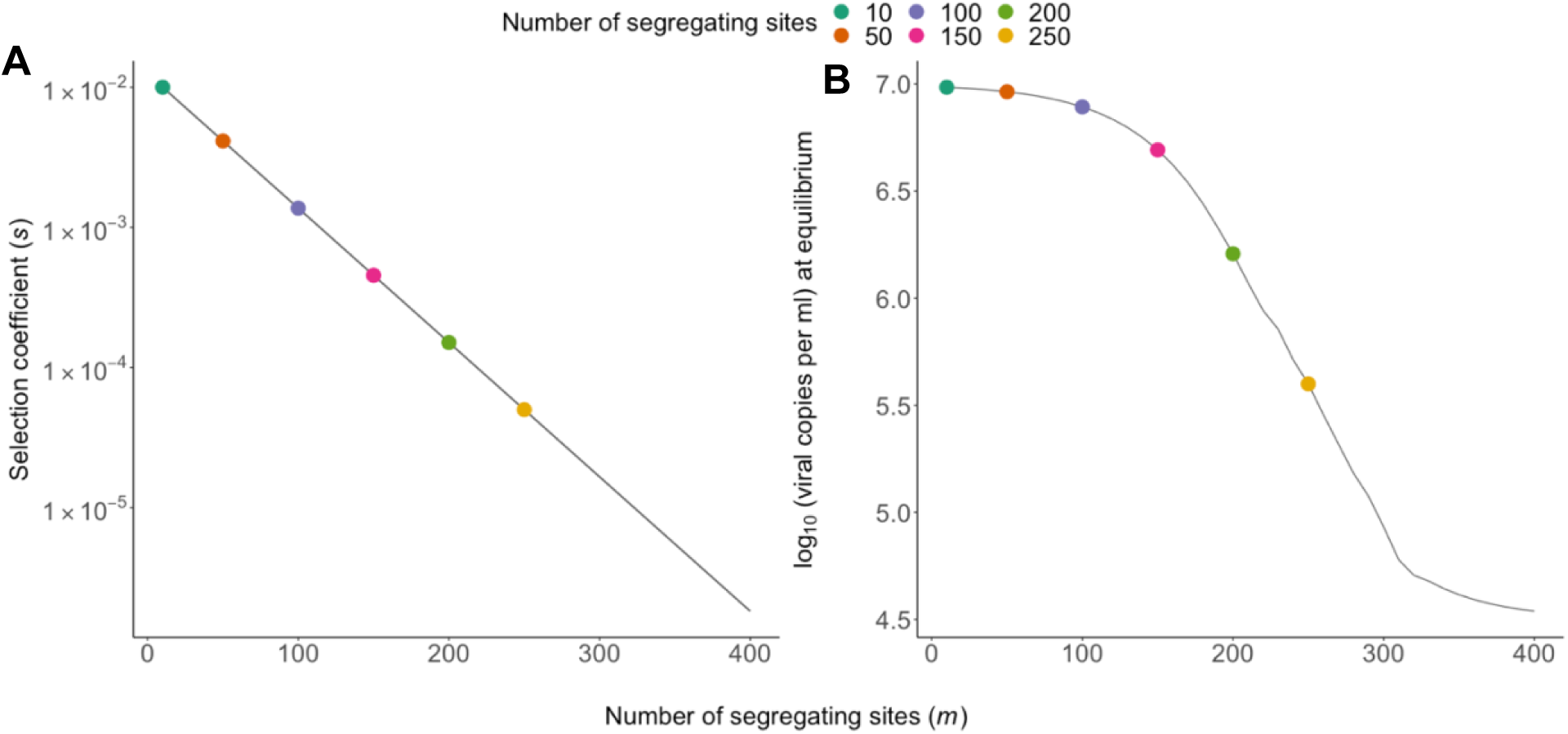
Within-host model fitness and equilibrium viral load. A) The relationship between the number of segregating sites and the fitness cost (synonymous with selection coefficient) associated with an additional mutation. We explore a range of values of *m*, with the corresponding fitness cost of a mutation reducing on a log-linear scale as *m* increases. The six choices of m explored here are indicated by coloured points. B) The viral load at equilibrium of an infection with an increasing number of segregating sites. As the number of sites increase and the fitness costs decrease, the population at equilibrium is characterised by mutation-selection balance that maintains deleterious viral types in the population, thus lowering viral loads.

With more segregating sites, the mutation-selection balance maintains a more diverse quasispecies, where multiple viral types coexist at equilibrium rather than a single dominant type (supp. Figure 1). Higher diversity in the within-host population creates a broader pool of variants available for transmission, which influences the trajectory of between-host evolution.

By adjusting how selection strength scales with the number of segregating sites, we could alter the model outcomes. If strong selection (high *s*) was maintained across more sites, equilibrium viral loads would be higher because deleterious mutations would be more efficiently removed. Conversely, if selection became weak at a lower number of sites, more deleterious mutations would accumulate, resulting in lower equilibrium viral load. While shifting the relationship between selection strength and number of sites would alter the specific equilibrium values, the qualitative behaviour of the system - and thus our main conclusions - would remain unchanged.

### Having many mutations with low fitness cost slows the tempo of within-host evolution

To fully explore the model and its dynamics, we considered five scenarios differing by the number of segregating sites (10, 50, 100, 150, 200, 250) and the fitness cost *s* of each mutation. The viral load calculation ensures that the virus type with the maximum number of mutations *(m*) has an associated viral load at the lower tail of reported spVLs, approximately 2 log_10_(viral copies per ml), and the fittest virus has an spVL of 7 log_10_(viral copies per ml). For each of the scenarios, we determined the evolutionary dynamics of the infection for all possible starting infection types. As expected, if there are few segregating sites, each with mutations of large effect, a within-host equilibrium was rapidly reached within months and was dominated by the fittest variants harbouring no mutations (Figure. 2A). Increasing the number of sites to 50 and 100 slows the within-host dynamics, however all infections have reached the maximum possible viral load within 5 years and 15 years respectively, with initially low vial loads steadily increasing throughout chronic infection.

**Figure 2.**
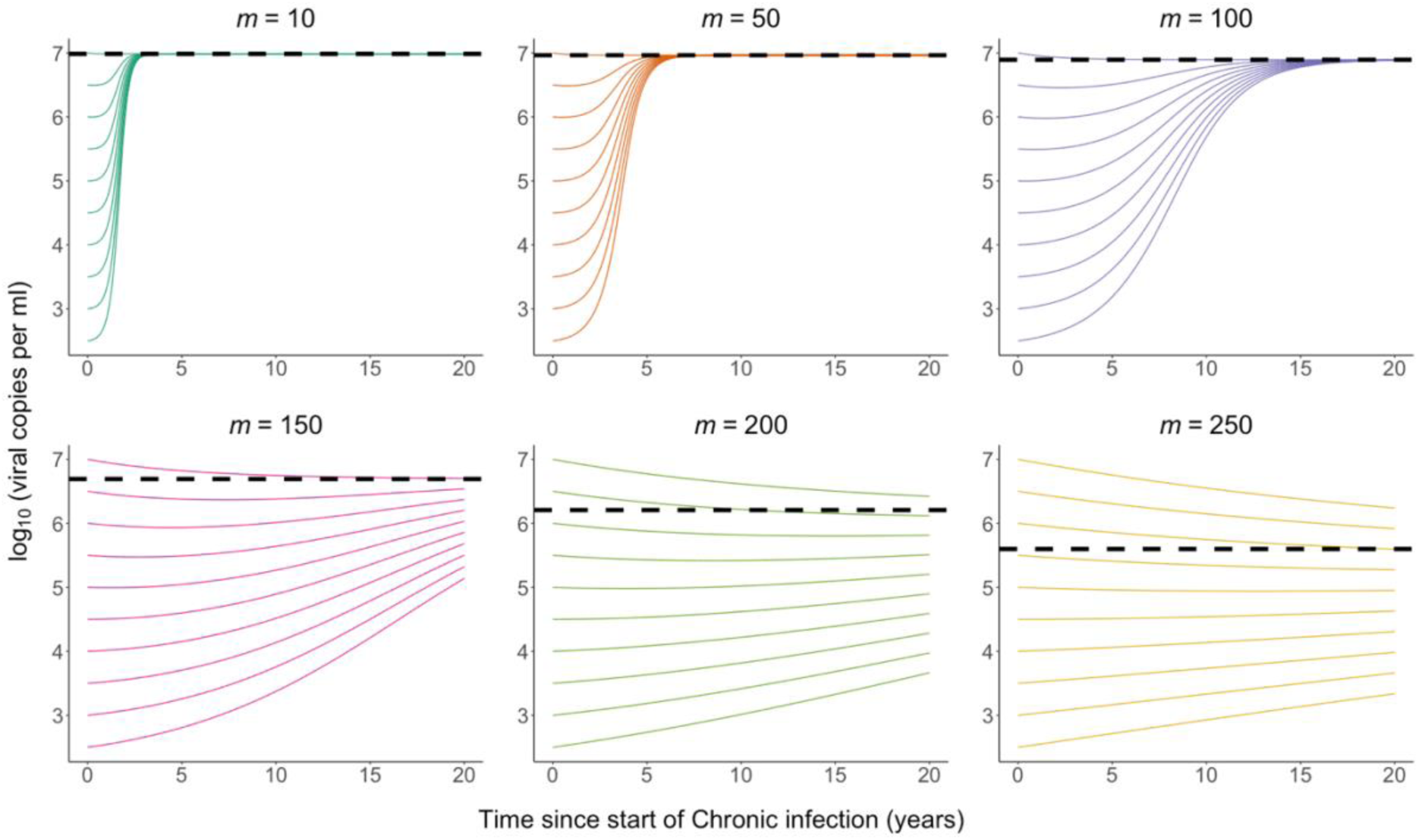
Within host viral load trajectories. The within-host viral load dynamics over time for 10, 50, 100, 150, 200 and 250 segregating sites, and varying initial numbers of mutations. The viral load of the equilibrium solution is shown by the black dashed horizontal line. The viral load changes as the virus population evolves over time, with new deleterious mutations appearing that are then lost due to purifying selection. When the cost of a mutation is high, viral loads climb to the maximum possible value. As we increase the number of segregating sites and reduce the fitness cost of a mutation, the selection against weaker virus types is balanced by the influx of mutations, and consequently the within-host dynamics are extremely slow relative to the typical duration of chronic infection.

When a large number (>100) sites are segregating and fitness costs are lower, a mutation-selection balance is reached at the within-host equilibrium state, lowering the viral load. Moreover, it took significantly longer than the assumed maximum infection duration to approach this equilibrium, depending on the number of mutations of the infecting strain. As we continue to increase the number of segregating sites from 200 to 250, the viral load dynamics do not qualitatively change within the time frame over which an infection typically occurs, and we continue to see stable viral loads.

### Many mutations with low fitness cost leads to between-host diversity in viral loads

To determine the expected distribution of spVLs among individuals predicted by our model, we first used numerical integration to determine the within-host dynamics of all possible *j*-type infections. For each infection-type, *j*, we could then calculate the mean viral load during chronic infection (our proxy for spVL), and in addition the infectivity of *i*-type virus at time *t* since infection, 𝛽_𝑖𝑗_ (𝑡) for all *i*. Using this information we determined the distribution of *j*-type infections, and therefore spVLs, at equilibrium by using a next-generation framework.

The distribution of spVLs varies substantially as we increase the number of segregating sites (Figure 3A-F). As the number of segregating sites increases, the average viral load at endemic equilibrium across individuals falls to the range of previously reported average spVLs, with 5.14, 4.78, and 4.562 (viral copies per ml) for 150, 200 and 250 segregating sites respectively (Figure 3D-F). The increased number of segregating sites leads to slower within-host dynamics and more diverse quasispecies populations, enabling the virus to evolve between hosts towards an intermediate spVL that maximises transmission potential. As a result, the epidemic size increases (Supp Fig 2).

**Figure 3:**
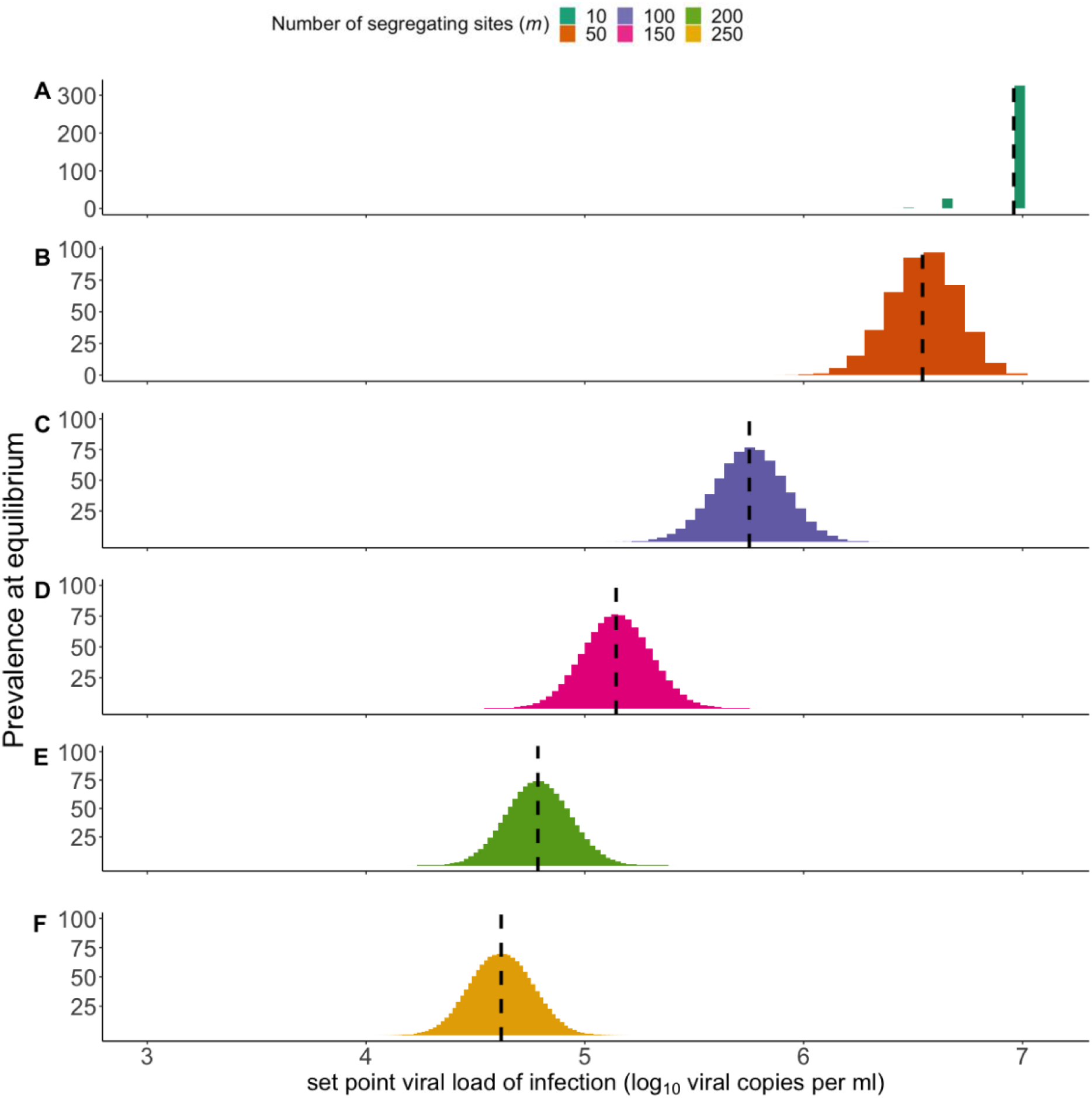
Between-host outcomes at endemic equilibrium. A) The distribution of set-point viral loads (spVL) at the between-host equilibrium. The fittest virus type rapidly dominates within-host and short-sighted evolution dominates. Black dashed lines denote the B-F) Greater within-host diversity provides a larger pool of variants that can be transmitted and on which between-host selection can act. As the number of segregating sites increases, the within-host dynamics slow and the virus is better able to evolve between-host towards an intermediate spVL that maximises transmission potential.

Whilst having many weakly deleterious mutations resulted in little variation in viral load over the course of an individual infection, the spVL during chronic infection differed by an order of magnitude between individuals. However, the variation in spVLs was still less than observed in infected populations (17). Host genetics are known to affect progression, and so we included host heterogeneity, such that the same infection type will induce different spVLs in individuals of different host type. For example, a host effect of 0.5 means that two individuals infected by the same virus type may differ in their spVL by a maximum of 1 log_10_ (viral copies per ml), 0.5 in either direction from the population mean. We find that in the case m=250, spVLs at the endemic equilibrium are more realistically diverse when a host-effect is assumed, and the number of circulating virus types increases, and the between-host diversity in spVL increases with host effect as expected (Figure. 4). We find the same effect on all choices of *m* (supp. Figure 3).

**Figure 4:**
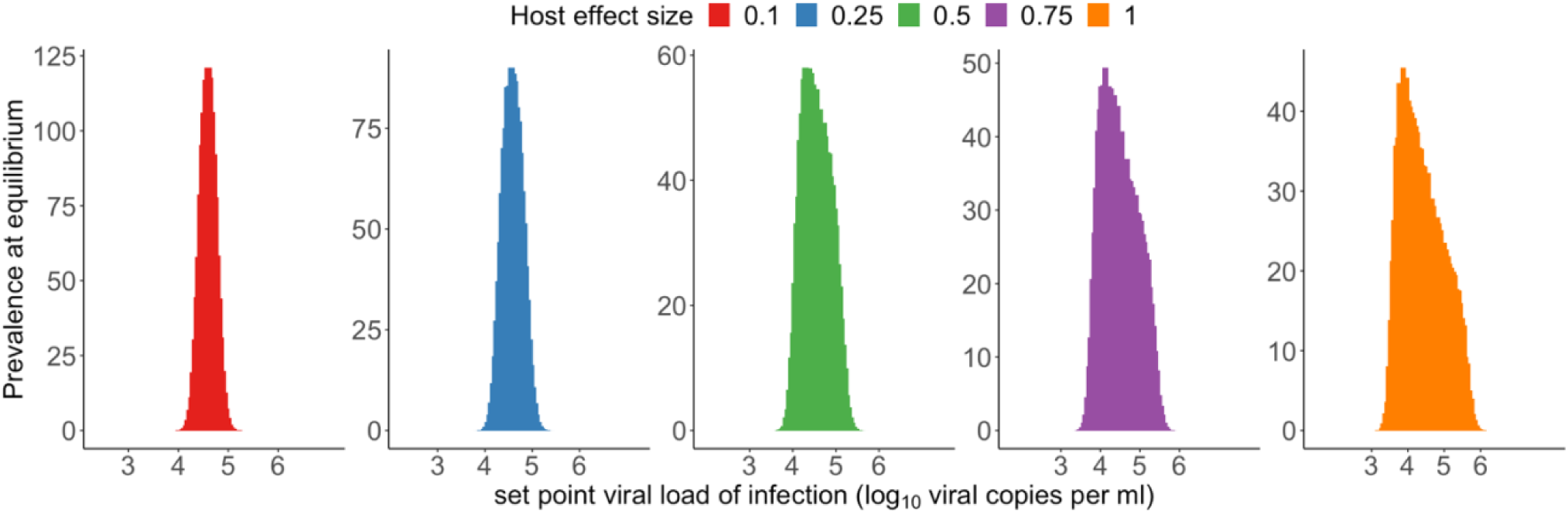
Between-host outcomes at equilibrium for an increasingly heterogenous host population. Histograms of spVLs in heterogenous infected population for different maximum host effect size, *e,* for 250 segregating sites. To account for the effect that host genetics has on viral load, we introduced a host specific additive effect to viral load. The size of the host effect is discretely uniformly distributed between -*e* and *e* and there are 50 host types. A maximum effect size of e=0.1 (A) results in a small increase in the range of viral loads observed, and as we increase *e* we observe a more realistic distribution of viral loads. Increasing the effect size towards *e*=1 further flattens and skews the distribution. Corresponding results for other choices of m are present in supp. Figure 4.

### Many mutations with low fitness cost results in viral load evolving to intermediate levels between-hosts

To capture the epidemiological dynamics and how the mean spVL varies as the epidemic progresses, we numerically integrated the full nested model over the first 100 years of an epidemic. For each scenario (choice of *m*), three simulations were run where the average spVLs of infection types at the start of the epidemic differed and initially there is a single infected individual. Population average viral loads decrease or increase depending upon the initial viral load, with cumulative population changes occurring over approximately a century for m=100 (fig. 5C), and over two centuries for larger m values (fig. 5D-E).

**Figure 5.**
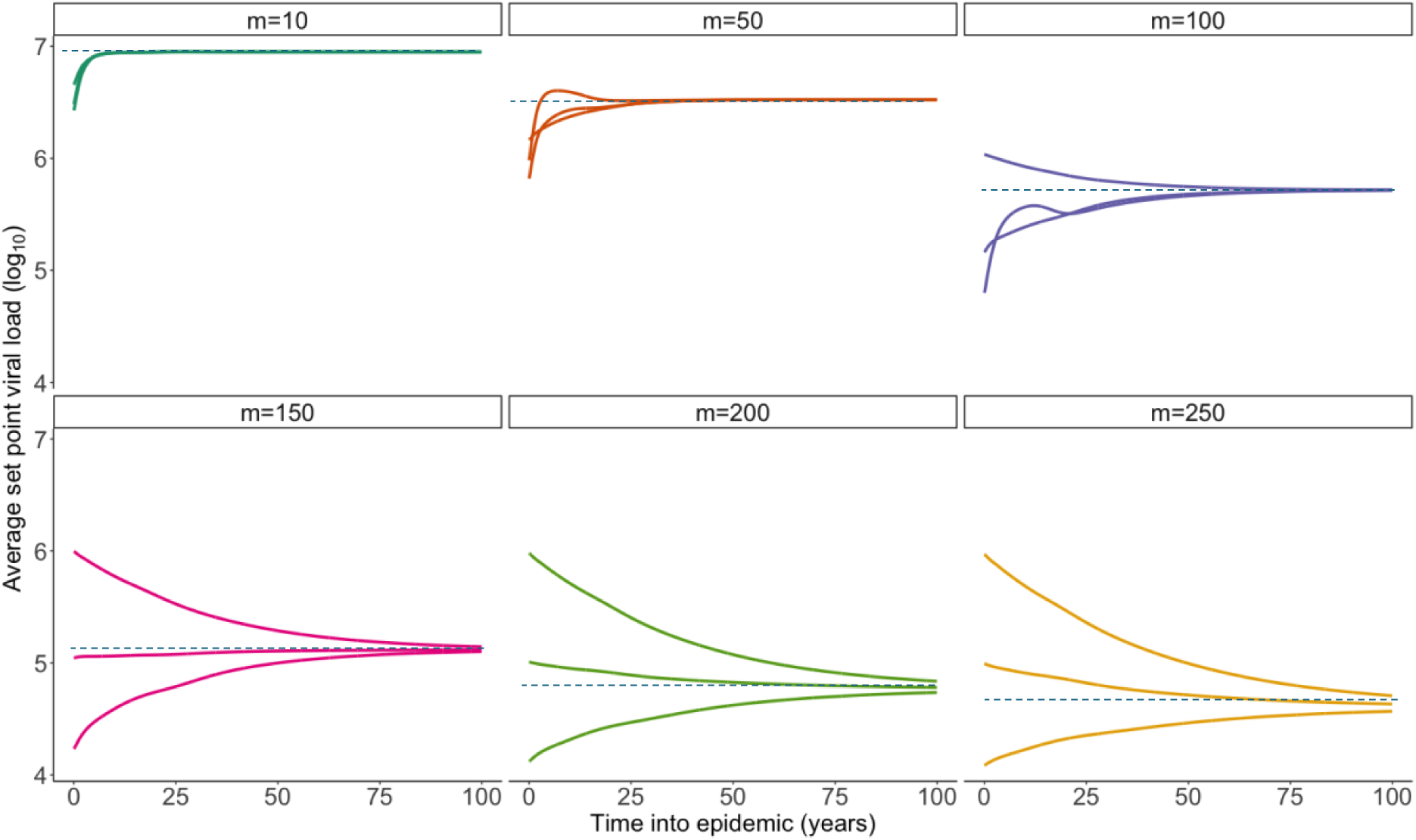
Average spVL over time in simulated epidemics. For *m*=10, short-sighted evolution leads to the rapid dominance of the fittest virus type as the single circulating variant within a few years of the epidemic. For *m*=50 and *m*=100, the within-host dynamics create a distinctive pattern at the population level. When epidemics start with lower viral load viruses that have a relatively long infection duration, the within-host evolution infections leads to the emergence and transmission of fitter variants with higher viral loads. This process accelerates the population-level increase in average spVL. When we assume a large number of weakly deleterious mutations (*m*=150,200,250) within-host dynamics are sufficiently slow for selection for transmission potential to influence the epidemiological dynamics and lower the average spVLs, despite the comparatively higher within-host equilibrium viral load. We observe slow cumulative changes in the average spVL, with over 100 years taken for convergence at higher values of *m*.

When we considered few mutations of larger effect, the fittest and most virulent variant rapidly outcompeted other virus types on the within-host scale. As a result, short-sighted evolution blocks between-host adaptation and the fittest variant dominates across the population at the expense of a reduction in transmission potential (fig. 5A- B). As the number of sites increase, the within-host dynamics slow and selection for transmission potential on the between-host scale drives the population-level dynamics, leading to gradual evolution towards an intermediate viral load. Importantly, we observe this dynamic for 100 segregating sites despite a high viral load at the point of mutation-selection balance. i.e. at within-host equilibrium. In this case, the fitness cost of each mutation is sufficiently small to slow within-host adaptation, resulting in stable viral loads during the first years of transmission opportunity.

### Viral loads are similar within transmission pairs

Within-host evolution can cause significant genetic change in the quasispecies between early infection and the time of onward transmission. In the absence of host heterogeneity or other environmental effects, we expect estimates of heritability to reduce in response to increased within-host evolution (11). We estimated heritability, ℎ^2^, by simulating transmission pairs in an infected population of homogeneous individuals at endemic equilibrium (Figure 6). Source infections were sampled based upon the infection-type population structure at the endemic steady state. Recipient infections were sampled based upon the transmission potential of each virus type during the infection of the source, and therefore the heritability estimates accounts for within-host evolution of viral factors.

**Figure 6:**
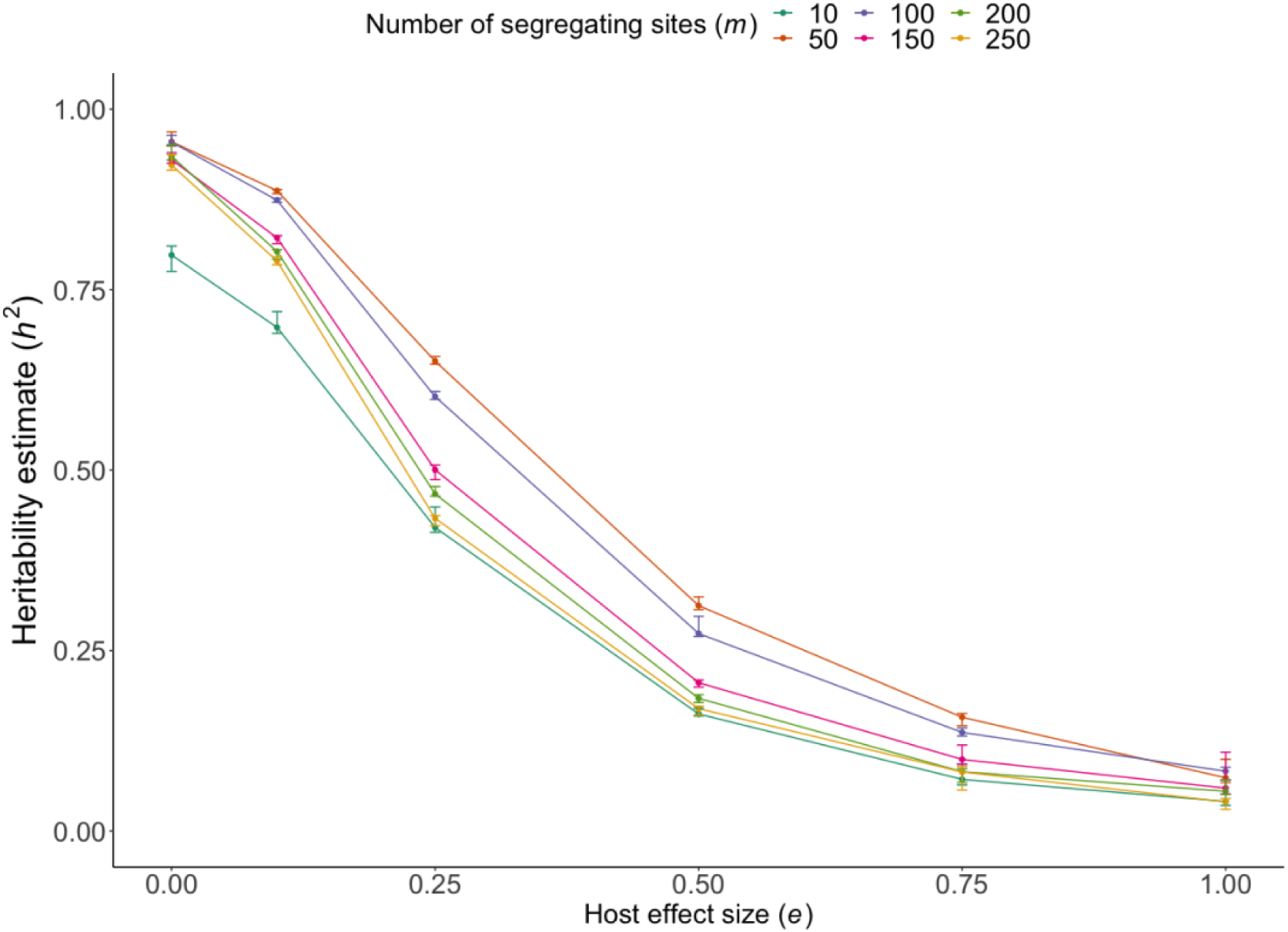
Heritability estimates. Heritability is estimated by a parent-offspring regression of spVL in simulated source and recipient pairs. The error bars indicate the standard deviation of the estimates taken from 1000 sampled sets of 500 transmission pairs. The infection type of the source (number of deleterious mutations of infecting virus type) is determined based upon the population-level prevalence of viral types at the endemic steady state. The virus type transmitted from source to recipient is determine based upon the transmission potential of each virus type over the course of the source infection. In a homogenous population, heritability is high, and naturally as a host-effect is introduced the amount of variability in spVL explained by the virus type, i.e. heritability, falls to within the range reported in multiple studies.

We find heritability is lowest for few mutations of large effect (*m*=10) due to rapid within-host evolution. In a homogenous population, all other scenarios have approximatley equal heritability, likely due to the initial period of stablity, high probability of transmission during acute infection, and the rapid changes occuring later into infection. As the host effect size increases, heritability decreases consistently across all scenarios, with the rate of decrease being similar regardless of the number of segregating sites, and heritability reduces to a range that has been reported in study estimates of broad-sense heritability. Heritability for *m*=50 and *m*=100 segregating sites is consistently greater than models with a larger number of segregating sites. This is because as the number of segregating sites increase, there is greater within-host variation in virus types, leading to more variation in transmitted viral loads between source and recipient.

## Discussion

There are several examples of mathematical models that predict pathogen dynamics by considering evolution across scales (38). With a multi-strain model that nests within-host dynamics of HIV within a between-host model, Lythgoe *et al*. (30) found that within-host evolution is rapid enough that HIV should evolve to high levels of virulence at the expense of fewer onward transmissions, and so within-host evolution was a greater prevailing force than evolution between hosts. By incorporating a reservoir into the model, evolutionary processes are delayed, and short-sighted evolution is prevented; however, the impact of the reservoir was highly sensitive to its assumed size (39). Van Dorp *et al*. (22) proposed an alternative model that considers immune escape and found that high host-heterogeneity in HLA types and consequently the escape and reversions of immune-escape mutations determine how spVL evolves. However, the model assumes that mutations are time-separated and occur according to a Markov process, effectively limiting the tempo of within-host evolution. Here, I applied the nested model framework from Lythgoe, Pellis and Fraser (30) without a latent reservoir and consider whether many weakly deleterious mutations can provide an additional and more parsimonious mechanism for spVL evolution under two levels of selection. If the number of segregating sites is sufficiently large, I show this can reconcile multiple paradoxes in HIV biology: the stability of viral loads during chronic infection despite high mutation rates, the heritability of spVL despite considerable within-host evolution between transmission events, and the evolution of spVLs that maximise transmission. This mechanism could also explain why heritable viral factors determining viral load have been so difficult to identify, due to the high statistical power needed to detect them.

Alternative solutions to the problem of rapid within-host evolution have also been proposed, including the existence of rugged and complex fitness landscapes that are difficult for the within-host viral population to traverse (30), the cycling of virus through the (unreplicating) HIV reservoir which then slows the rate of within-host evolution (39) and viral factors that are heritable but are not under within-host selection, specifically polymorphisms that target the cell activation rate and are therefore beneficial to the entire virus population (40). It is, however, difficult to reconcile the latter theory with the relationship between viral load and replicative capacity, and none of the described theories provide a fully satisfactory explanation of a heritable set-point viral load that varies by orders of magnitude between individuals, or why viral virulence factors have been so hard to identify (19).

We do not expect this mechanism alone to fully control viral load, as host factors— such as protective HLA alleles—are well-documented contributors to viraemic control (18,41–43). These host factors shape the immune system’s ability to suppress viral replication and explain part of the variation in viral load across individuals. Specific viral mutations with large effects on fitness have been identified (44), but these are relatively low in number and the total narrow-sense heritability from all genetic hits is significantly lower than our estimates of broad-sense heritability. It is therefore likely that viral genetic factors each carry a small effect that studies do not have the statistical power to detect. Virus-host interactions further complicate this picture. The complex relationship between viral evolution and host immune responses creates a dynamic environment where the effects of individual genetic variants may be context-dependent, making it challenging to identify robust GWAS hits associated with spVL.

HLA footprints have been associated with spVL heritability, as viruses adapted to the source’s HLA profile maintain higher viral loads when transmitted to recipients with similar HLA backgrounds (29). However, this mechanism cannot explain the dominant power of between-host evolution and the selection for maximal transmission fitness at a population level (22).

Studies of the HIV fitness landscape have identified the significant contribution from mutations that carry a very small fitness cost to replicative capacity, with these largely being synonymous (45). Synonymous mutations, while not directly affecting protein structure, can still impede viral fitness by influencing RNA stability, translation efficiency, or protein folding. The deleterious component of the landscape has been shown to be universal across infections and fundamental to the high diversity within HIV group M diversity (36). Further support for our model may also be found by investigating testable predictions with the appropriate datasets. First, we would expect a correlation between the number of weakly deleterious mutations in a within-host virus population and viral load. We therefore can consider rare alleles and their impact on viral load in cohorts of individuals where viral load and viral sequencing data is available.

As with all mathematical models, the model proposed here represents a significant simplification of complex biological processes for the sake of tractability and interpretation. Though this model reproduced known behaviour, in reality other processes will contribute to varying degrees, such as repeated immune escape and reversion across different host environments, and perhaps a small number of mutations of large effect. Developing an informed understanding of the virus factors that control virulence, and how they evolve in response to selection at the within- and between-host scales, will ultimately provide important insights into the severity of viral infections, including HIV, how this might change through time, and improve future treatments and public health policy.

## Code Availability

The scripts to perform the analysis can be found at the GitHub repository https://github.com/HLongleyOx/EvolutionOfVirulence2025

## Acknowledgments

H. Longley is supported by the Engineering and Physical Sciences Research Council Centre for Doctoral Training in Health Data Science (EP/S02428X/1). K. Lythgoe is supported by the Royal Society and the Wellcome Trust (107652/Z/15/Z) and the Li Ka Shing Foundation. C. Fraser is supported by the Li Ka Shing Foundation.

---

**Figure 1.**
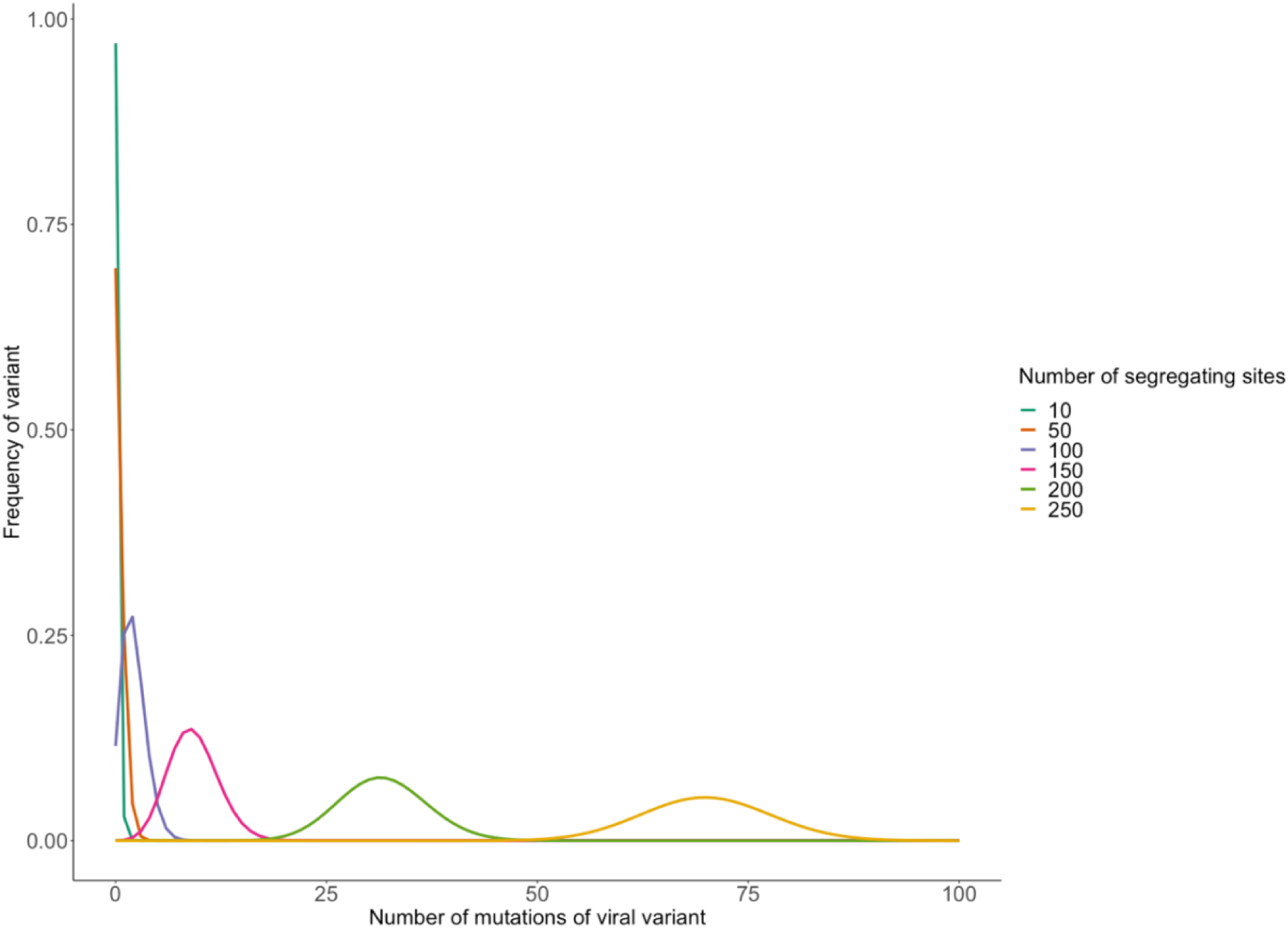
The frequency distribution of the viral population at equilibria. A specific viral type is defined by its number of mutations. When we consider few mutations of large effect, the population is dominated by a single virus type of high relative fitness. As we increase the number of segregating sites and lower the associated fitness cost, the population becomes increasingly diverse, which has implications for the viral variants that are transmitted and between-host evolution. This diversity in the within-host viral population creates a broader pool of viral variants available for transmission, potentially influencing both transmission dynamics and the trajectory of between-host evolution

**Figure 2.**
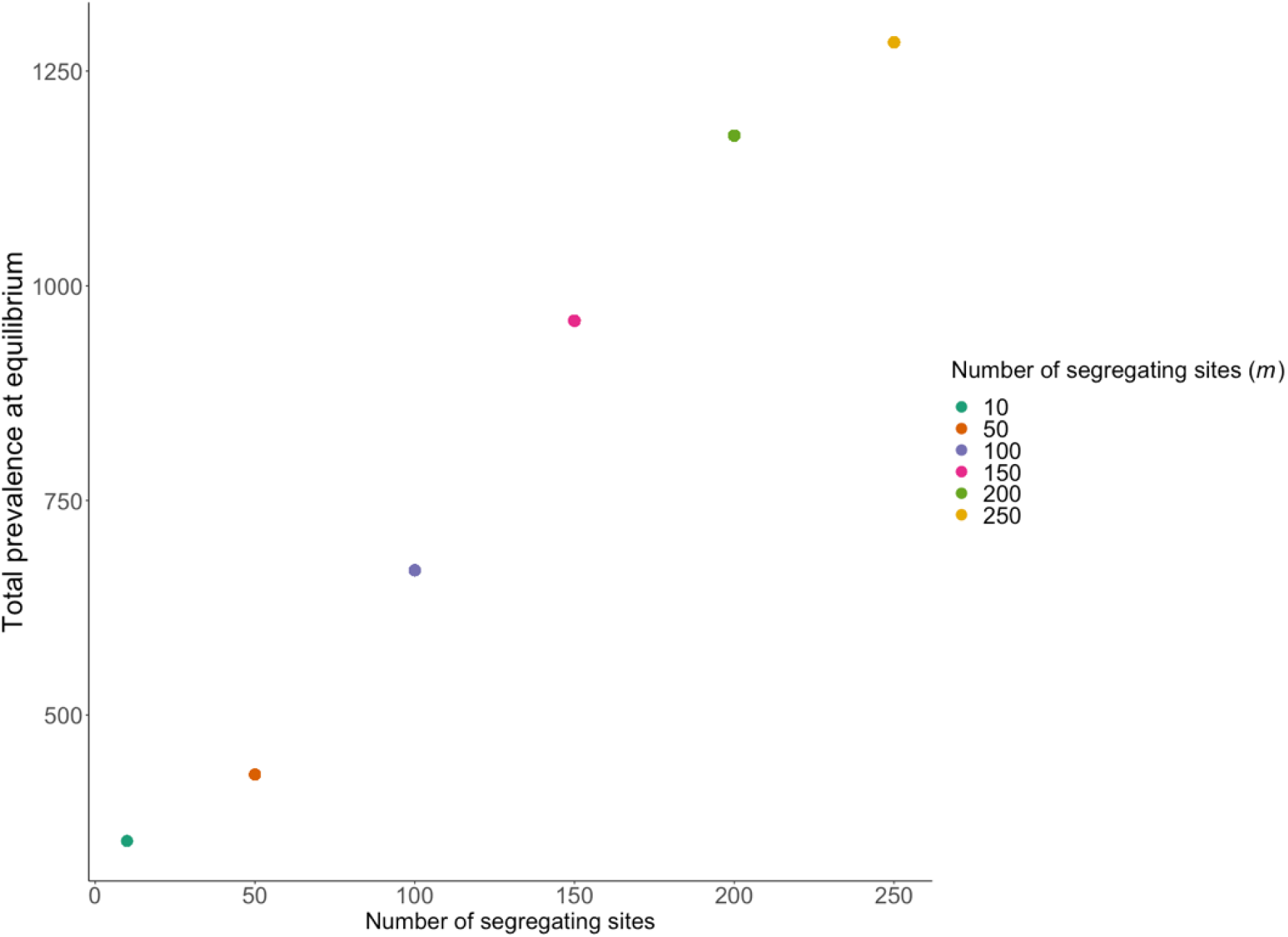
The total prevalence at the endemic steady state. As the number of segregating sites increases, the within-host dynamics slow down and the viral population is more diverse. As a result, between-host selection is able to select the virus types with greatest transmission potential, ultimately increasing the endemic prevalence.

**Figure 3.**
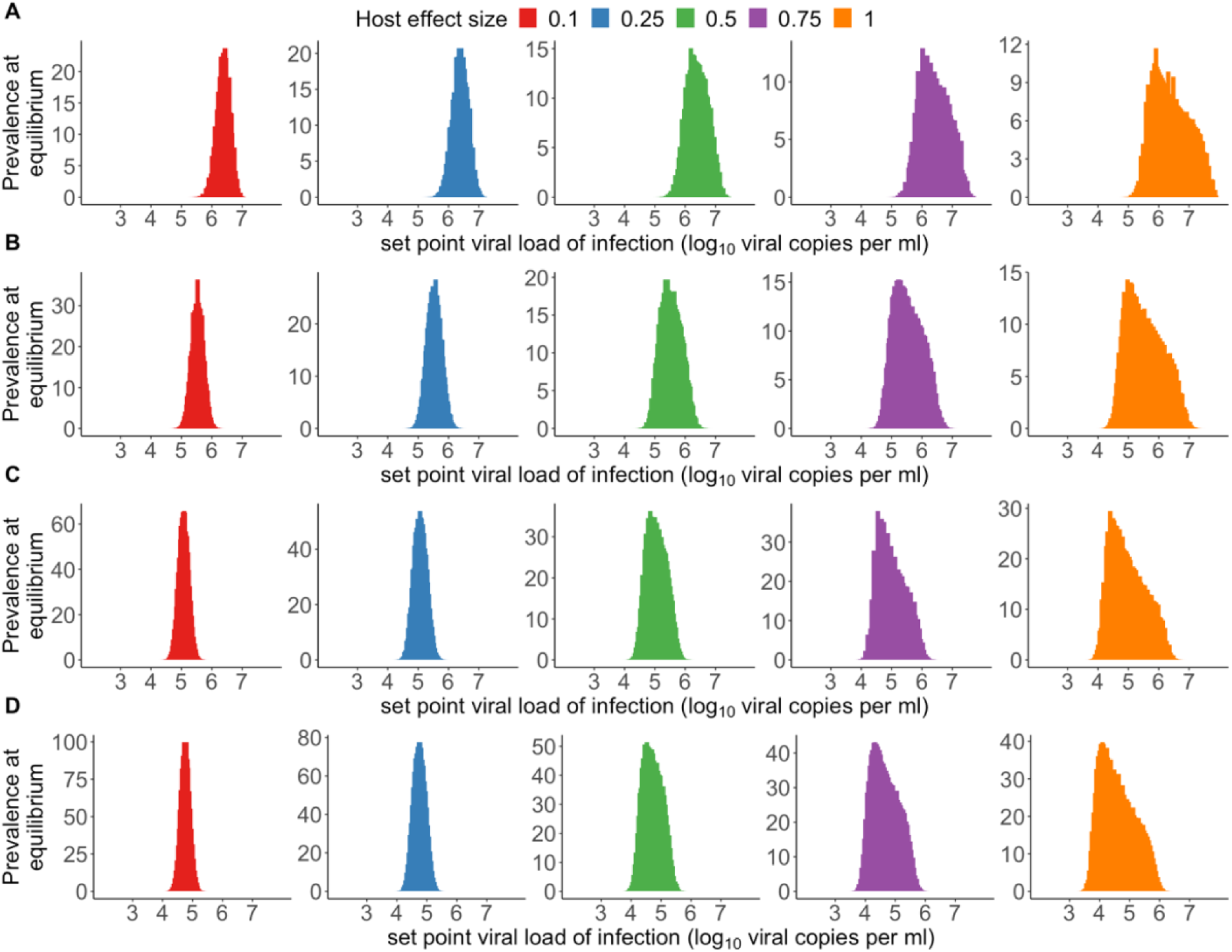
Between-host outcomes at equilibrium for an increasingly heterogenous host population for all models. Histograms of spVLs in heterogenous infected population for different maximum host effect size, *e,* for A) 50 B) 100, C) 150 and D) 200 segregating sites. To account for the effect that host genetics has on viral load, we introduced a host specific additive effect to viral load. The size of the host effect is discretely uniformly distributed between -*e* and *e* and there are 50 host types. Increasing the host-effect broadens the distribution of spVLs.

